# Cocaine sensitization in male rats requires activation of estrogen receptors

**DOI:** 10.1101/2024.02.07.579327

**Authors:** Raissa Menéndez-Delmestre, José L. Agosto-Rivera, Amanda J González-Segarra, Annabell C. Segarra

## Abstract

Gonadal steroids play a modulatory role in cocaine use disorders, and are responsible for many sex differences observed in the behavioral response to cocaine. In females, it is well established that estradiol enhances the behavioral response to cocaine. In males, we have recently shown that testosterone enhances sensitization to cocaine but its mechanism of action remains to be elucidated. The current study investigated the contribution of DHT, a non-aromatizable androgen, and of estradiol, in regulating cocaine-induced sensitization in male rats. Gonadectomized (GDX) male rats treated with estradiol sensitized to repeated cocaine administration, while GDX rats treated with DHT did not, implicating estradiol in cocaine sensitization. Furthermore, intact male rats treated with the antiestrogen ICI 182,780 did not show sensitization to repeated cocaine. This study demonstrates the pivotal role of estradiol in cocaine-induced neuroplasticity and neuroadaptations in the rodent brain.

## Introduction

Gonadal steroids play a major role in the establishment and maintenance of sexually dimorphic brain structures and behaviors, such as drug addiction (Segarra, 1993; Segarra et al., 2010; Carr et al, 2020). Although some of these differences may be attributed to the diversity of social and environmental surroundings, fundamental biological differences exist between the sexes that contribute to psychological and physiological differences in addictive behaviors.

Animal studies find that females learn to self-administer cocaine faster, their intake is greater (Lynch and Carroll, 1999; Yang et al., 2007, Algallal et a., 2019), they require less training sessions (Russo et al., 2003), their locomotor response to cocaine is higher (Forgie and Stewart, 1993; Harrod et al., 2005a; Hu and Becker, 2008; Segarra et al., 2010; Bobzean et al., 2014) and they develop conditioned place preference to cocaine at lower doses (Russo et al., 2003; Zakharova et al., 2009) than males. Not all studies find a difference between the sexes or agree in the differences found (Caine et al., 2004; Algallal et al, 2019; Kawa and Robinson, 2019). These discrepancies between studies may be attributed to methodological differences, such as the dose of cocaine administered, the schedule of reinforcement in self-administration studies, the behavior assayed (training vs acquisition), the parameters measured and prior drug exposure history among others.

The mechanisms that mediate this differential response to cocaine have not been fully elucidated. Studies investigating sex differences in cocaine pharmacokinetics have yielded conflicting reports. Whereas many report no sex differences (Bowman et al., 1999; Mendelson et al., 1999; Evans and Foltin, 2010; Coe et al., 2018, McDougall et al., 2018) others find the metabolism of cocaine varies with sex (Niyomachi et al., 2066; Visalli et al., 2005). It must be emphasized that pharmacokinetics studies are complex, and multiple factors, such as age, dose, individual differences in metabolism among others can affect the results obtained. One factor that has received more attention is the role of gonadal steroids in mediating these different behavioral responses. For example, in females estradiol enhances cocaine-induced sensitization (Segarra et al., 2014; Souza et al., 2014), conditioned place preference (Segarra et al., 2014) cocaine self-administration (Hu and Becker, 2008; Lynch et al., 2002; Perry et al., 2013; Ramôa et al., 2013) and the motivation to seek drugs

The role of gonadal steroids in regulating addictive-like behaviors in males is not clear. In mammals, plasma gonadal steroids are synthesized mainly by the gonads and to a lesser extent, by the adrenal cortex. However, neurons and glial cells also synthesize gonadal steroids *de novo* and these contribute significantly to their neuroprotective effects on neural tissue (Hojo et al., 2004; Jellinck et al., 2007). In males, the main circulating gonadal steroid is testosterone. Testosterone is also a prohormone, and serves as a substrate for the production of other steroids, such as dihydrotestosterone (DHT). The production of DHT is dependent on the presence of the enzyme 5-α reductase in target tissue (Pelletier et al., 1994). Testosterone and DHT bind to the cytoplasmic androgen receptor (AR) widely distributed in skeletal muscle, male reproductive tract and neural tissue, although DHT has a higher affinity for the AR than testosterone. When these hormones bind to the AR they elicit several changes and cause translocation of the hormone-receptor complex to the nucleus. There it associates with coregulators that affect AR stability, binding to androgen response elements in the DNA, and chromatin remodeling among others (for review see Bennet et al., 2010; Solomon et al., 2019).

Testosterone can also be converted to estradiol, the main gonadal steroid in females, by the cytochrome P450 aromatase (CYP19), found abundantly in brain tissue (Roselli et al., 1998; Roselli and Resko, 1987). Estradiol binds to a different family of receptors, the estrogen receptor (ER). These hormone receptors are present in several brain regions of males and females such as those that regulate addictive behaviors, (González et al., 2007; Mitra et al., 2003; Pelletier, 2000). Thus, testosterone can exert its effects on brain tissue by binding to ARs or indirectly, by activating ERs.

Our recent studies show that testosterone modulates the locomotor response to cocaine. GDX male rats show greater cocaine-induced hyperactivity to a single cocaine injection than intact males or GDX males that received testosterone replacement (Menéndez-Delmestre and Segarra, 2011). In contrast, the locomotor response to repeated cocaine administration increases over time only in gonadally intact rats, or in GDX rats that received testosterone, indicating that testosterone is necessary for male rats to display cocaine-induced behavioral sensitization (Menéndez-Delmestre and Segarra, 2011).

Behavioral sensitization is defined as the successive augmentation of locomotor hyperactivity elicited by repeated administration of a drug, such as cocaine. The psychomotor stimulant effects are mediated, at least partially, by the neural substrates that regulate its reinforcing properties as well, such as the mesocorticolimbic dopaminergic system (Wise and Bozarth, 1987). It has been argued that the addictive liability of a drug can be predicted by its ability to induce psychomotor sensitization (Wise and Bozarth, 1987). As such, sensitization involves neuroadaptations in the mesocorticolimbic system that contribute to changes in the motivational circuitry underlying craving and relapse (Chefer et al., 2005; Kumar et al., 2005; Thomas et al., 2008). Although it has several limitations, it induces measurable long-lasting changes in mesocorticolimbic substrates (Kuhn et al., 2020, Carr et. al., 2020).

Similar to our findings, others studies report that gonadectomy enhances the locomotor response to a single injection of cocaine (Beatty et al., 1982; Menéndez-Delmestre and Segarra, 2011; Sorg et al., 2002; Walker et al., 2001) or of amphetamine (Dluzen et al., 1986). When testosterone was administered to intact or to GDX males, it attenuated amphetamine (Beatty et al., 1982; Forgie and Stewart, 1994) and cocaine-induced locomotor activity (Chen et al., 2003; Long et al., 1994; Menéndez-Delmestre and Segarra, 2011; Sorg et al., 2002; Van Luijtelaar et al., 1996) as well as cocaine self-administration (Mello et al., 2011). Nonetheless, there are studies that fail to find an effect of gonadectomy or of testosterone replacement (Caine et al., 2004; Chin et al., 2002; Haney et al., 1994; Harrod et al., 2005a, 2005b; Hu and Becker, 2003; Jackson et al., 2006; Menniti and Baum, 1981; Minerly et al., 2008). Methodological differences may account for some of the differences in the results obtained. For example, in one study testosterone propionate (2 mg/kg) is injected 30 min before psychostimulant administration (Chen et al., 2003), limiting many of the effects that may be genomic-dependent. In addition, the gonadal state of the animals varied, some studies administered testosterone to gonadally intact animals, in other studies animals were gonadectomized. Furthermore, the dose, form of administration (injected or via subcutaneous Silastic tubes or pellets) and chemical form of testosterone (propionate, acetate), which affects the half-life and binding properties, is not consistent among studies (Caine et al., 2004; Chen et al., 2003; Long et al., 2000; Menéndez-Delmestre and Segarra, 2011; Minerly et al., 2010).

Our goal was to determine the hormonal signal by which testosterone potentiates the behavioral response to cocaine over time. Given that testosterone interacts directly with ARs, we hypothesized that testosterone’s effects on cocaine sensitization in males might be mediated through activation of the androgen receptors. We predicted that GDX male rats treated with DHT, a non-aromatizable androgen, would show enhanced sensitization. The alternative was that testosterone was converted to estradiol and interacted with ERs. To test these hypotheses, male rats were gonadectomized, and received an empty implant (GDX), an implant with DHT (GDX-DHT) or an implant with estradiol benzoate (GDX-EB). To further explore the role of ERs in cocaine-induced sensitization in male rats, gonadally intact male rats were treated with the antiestrogen ICI 182,780 intracerebroventricularly to determine if central ERs participate in the development and/or expression of cocaine-induced sensitization of male rats.

## Materials and Methods

### Animal Subjects

Adult male Sprague-Dawley rats (320g) were purchased from Taconic Farms and housed in pairs in a temperature and humidity-controlled room. They were kept in a 12-hr light-dark cycle (lights off at 7 pm) with water and Harlan Tek™ rat chow provided *ad libitum*. Animals were allowed to acclimate to the Animal Facilities for 5 days before any experimental manipulation. The experiments required testing of several groups at the same time (i.e. GDX (Saline and Cocaine), GDX-EB (Saline and Cocaine) Intact-Sham (Saline and Cocaine). To achieve this, and have an adequate number of animals per group, experiments were repeated several times (12 in the case of GDX and Intact-Sham rats).T he figures prepared for this manuscript represent a pool of all these experiments (See Table 1 of Appendix). All animal experimental procedures were approved by the Institutional Animal Care and Use Committee (IACUC) of the University of Puerto Rico, Medical Sciences Campus and adhere to USDA, NIH and AAALAC guidelines.

### Gonadal Steroid Implant preparation

#### Silastic implants

To minimize the stress of injections and the fluctuations in plasma levels of hormones associated with bolus steroid injections, hormones were replaced using Silastic implants. These are prepared according to the method described by Legan et al.(1975), and modified according to Febo et al.(2002). The Silastic tubes (Dow Corning, Midland, MI, USA) had an internal diameter of 1.47 mm and an external diameter of 1.97 mm (“membrane thickness” of 0.5 mm). For estradiol administration, the length of the Silastic tubes was 5 mm (volume = 8.49 mm^3^). For DHT administration we used 2 Silastic implants, each measuring 10 mm in length (total volume = 16.97 mm^3^). Implants were empty (GDX) or filled with gonadal steroids and sealed on both ends with Room Temperature Vulcanizing (RTV) Silicone Sealant (Dow Corning, Midland, MI, USA). Hormones were purchased from Sigma-Aldrich (St. Louis, MO, USA). Implants were placed subcutaneously in the midscapular region during surgery and remained there throughout the experiments. Silastic implants were used instead of commercially available steroid pellets. Our previous studies agree with that of others that report a better and more consistent release from Silastic tubes (Strom et al. 2008; Ingberg et al. 2012) than from commercially available pellets.

#### Estimated estradiol values

- The Silastic implants were filled with approximately 4 mg of estradiol benzoate (EB). Taking into consideration the values reported in the literature, our calculations indicate that **the range of estradiol achieved by our implants should be between 50 and 200 pg/ml.** These were calculated using the values obtained in males by DeVries et al. (1986) as the lower end and those obtained by our lab in females (Mosquera et al, 2015) as the higher end (See Table 1 in Appendix).

#### Estimated DHT values

- The Silastic implants were filled with approximately 10 mg of dihydrotestosterone benzoate (5α-androstan-17β-ol-3-one benzoate, DHT). According to **our calculations, the range of DHT achieved by our implants in males should be between 1.2 - 90 ng/ml.** These were calculated using the values obtained by Menniti et al. (1980) as the lower end and those of De Vries et al., (1986) as the higher end.

### Surgical procedures

#### Orchidectomy

Following the acclimation period, a group of animals was orchidectomized using ketamine (65 mg/kg) and xylazine (7 mg/kg) (Sigma-Aldrich, St. Louis, MO, USA) injected intraperitoneally as anesthetics. Intact animals were sham operated and remained gonadally intact (INT-Sham). After 7 days of recovery, animals were tested for behavioral sensitization.

#### Stereotaxic surgeries: Administration of ICI-182,780

ICI-182, 780 (also known as fulvestrant and Faslodex®) is a highly selective antiestrogen (Howell et al., 2000) that impairs dimerization of ER, increases ER turnover, disrupts nuclear localization, and downregulates the ER (Dauvois et al., 1993; Howell et al., 2000; Parker et al., 1993). ICI 182,780 effectively blocks estradiol’s mitogenic effect on the uterus, and attenuates estradiol’s effect on growth and sexual receptivity (Wade et al., 1993). Therefore, to investigate if centrally located ER promote cocaine-induced locomotor activity in males, we administered ICI-182,780 to gonadally intact male rats (INT+Veh) and assessed the stimulant effects of cocaine using the behavioral sensitization paradigm. ICI-182, 780 (Sigma-Aldrich), was dissolved in 22.5% of (2-hydroxypropyl)-β-cyclodextrin and 0.45% saline solution (vehicle) and administered intracerebroventricularly (icv) via an Alzet mini osmotic pump for constant administration of ICI-182,780 or vehicle (0.15 ul/hour). The concentration delivered of ICI-182,780 was 0.075ug/hour. The Alzet pumps delivers drug (and vehicle) for 28 days (according to the manufacturers specifications).

Gonadally intact males (INT+Veh) were anesthetized with 50 mg/kg of sodium pentobarbital (i.p.) and a cannula was inserted into the right lateral ventricle (coordinates: AP=-0.72mm, ML= +1.6mm, DV=-4.0mm from Bregma). The cannula was attached to an osmotic mini-pump (ALZET model 2006 and ALZET Brain Infusion Kit) that was placed subcutaneously near the scapular region; the rate of delivery was 0.15uL/hour. One group of animals received osmotic mini-pumps filled with vehicle (VEH group) and the other group received osmotic mini-pumps with ICI-182, 780 (ICI group). After 7 days of recovery, all animals were submitted to behavioral testing. At the end of the experiment, animals were euthanized and the accuracy of the lateral ventricle cannula was verified histologically. Animals with incorrect cannula placement were not included for data analysis.

### Behavioral Tests

#### Behavioral Apparatus

Horizontal and stereotyped activity was measured using 10 automated animal activity cage systems (Versamax^TM^ system) purchased from AccuScan^TM^ Instruments (Columbus, Ohio, USA). The activity cages were made from clear acrylic (42 cm x 42 cm x 30 cm), with 16 equally spaced (2.5 cm) infrared beams across the length and width of the cage at a height of 2 cm from the cage floor (horizontal beams). The cage contained an additional set of 16 infrared beams at a height of 10 cm from the cage floor (vertical beams). All beams were connected to a Data Analyser^R^ that sent information to a personal computer that displayed beam data through a Windows^R^-based program (Versadat^R^).The Versamax^TM^ system differentiates between stereotyped and horizontal locomotor activity based on repeated interruption of the same beam or sequential breaking of different beams, respectively

#### Cocaine-induced locomotor activity

The locomotor response to cocaine was measured in a dimly lit isolated room (below 50 lux) with constant temperature (25°C) and humidity. To diminish the effects of novelty, animals were habituated to the activity cages for one hour (Day 0) a day before receiving the first saline or cocaine injection. From days 1-5, animals received daily i.p. injections of 0.9% sterile saline (Nicolas Carrillo, Baxter Healthcare Corporation, Deerfield, IL, USA) or of cocaine (15mg/kg, Sigma-Aldrich, St Louis, MO, USA). During days 6-12 animals remained in their home cages undisturbed. On day 13, animals were challenged with an injection of saline or cocaine (15mg/kg). Locomotor activity (LMA) was recorded on days 1, 5 and 13 for 90 mins. The first 30 min, animals were placed in the activity cage to reduce novelty-induced activity and to determine basal locomotor activity (LMA). Animals were then injected with saline or cocaine and LMA recorded for an additional 60 min immediately after injection (Fig 1). Each behavioral testing session included at least one animal of each group to minimize intergroup variation that may result from differences in time of testing and/or injections. The LMA of each group on day 1 was compared to that of day 5 and of day 13. A significant increase in LMA after repeated cocaine administration (i.e., day 1 vs day 5) indicated sensitization. After the last behavioral test (Day 13), animals were euthanized.

**Figure 1.**
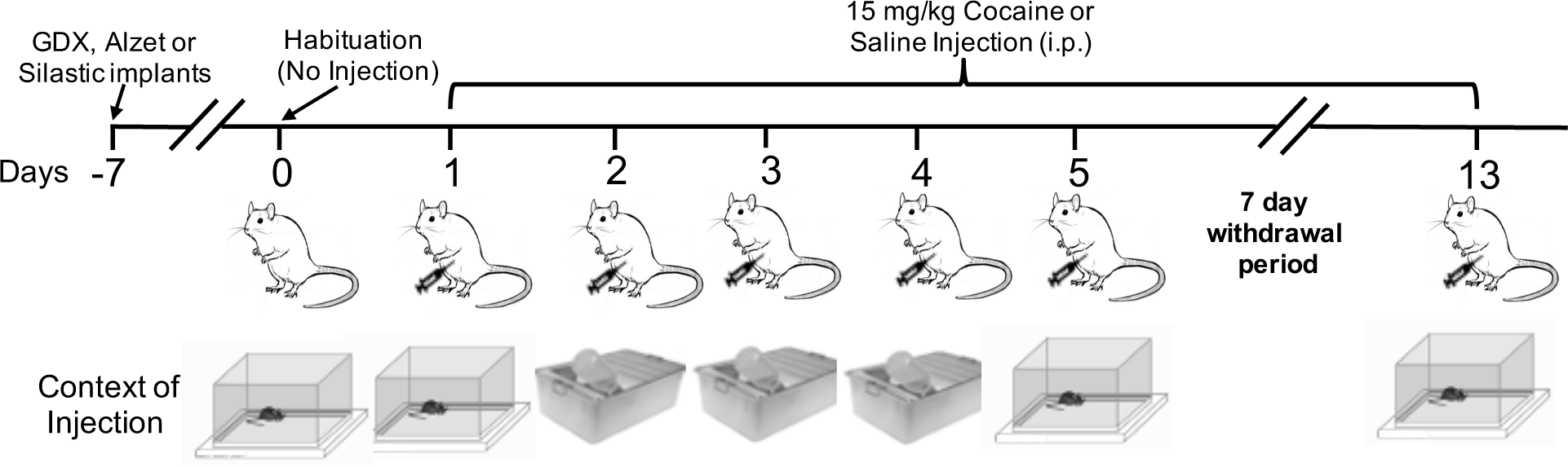
Behavioral sensitization protocol. Five days after arrival to the Animal Facilities, rats were gonadectomized (or sham gonadectomized) and received a Silastic implant (Day −7). The INT-Veh and INT-ICI groups were also subjected to stereotaxic surgery and received an Alzet™ osmotic mini-pump. All animals were allowed a 7 day recovery period (Day −7 to Day 0). On Day 0, rats were habituated to the locomotor activity chamber for 1 hour. From Days 1 through 5 and on Day 13, rats were injected with saline or with cocaine (15 mg/kg). On days 1, 5 and 13 rats were injected in the locomotor activity chamber, on days 2, 3 and 4 animals were injected in their home cages (see context of injection). From days 6 through 12, rats remained undisturbed in their home cages.

### Statistical Analysis

All data were analyzed using GraphPad (GraphPad Software, San Diego California USA). Data were analyzed using area under the curve (AUC) of the plotted horizontal activity. We selected AUC because the data distribution in all groups followed a Gaussian distribution without the need of transformations. In addition, the AUC measure was more sensitive in detecting changes in locomotor activity among the groups. We selected minutes 30 to 50 of the time course since the locomotor activating effects of cocaine are greatest during the first 20 minutes after injection.

An unpaired Student t-test was used to compare the acute locomotor response to cocaine (Day 1) of two groups and a One-Way ANOVA was used when comparing more than two groups (Table 4 of Appendix). To ascertain if rats showed sensitization, the timecourse of each group was analyzed separately using a RM ANOVA with days (1, 5 and 13) and minutes (35-90) as the repeated factors. Tukey’s multiple comparisons Test was used as post-hoc analysis to compare each timepoint (Table 5 of Appendix). To determine differences in sensitization, the percent change of days 5 and 13 from that of day 1, was calculated for each group as previously described (Segarra et al, 2014). The groups were compared using a Two-Way Repeated Measures ANOVA (Two-Way RM ANOVA), using “Days” as the repeated measure and “Treatment” as the independent variable followed by a post-hoc test (Table 4 of Appendix). Data are presented as the mean ± standard error of the mean (SEM). A p-value less than 0.05 (p<0.05) was considered statistically significant.

## Results

### Orchidectomy increases cocaine hyperactivity in male rats

Male rats were gonadectomized and tested for behavioral sensitization to cocaine. Confirming our previous results (Menéndez-Delmestre and Segarra, 2011), gonadectomy increased the acute locomotor response to cocaine (Fig. 2C; Unpaired Student T-test, t=2.988, df=58.07, p=0.0041). In addition, an increase in cocaine-induced hyperactivity was observed in INT-Sham (Fig 2A: Two-Way RM ANOVA, Days, F_(1.9, 71.5)_ = 12.59; p<0.0001) and to a lesser extent, in GDX male rats (Fig 2B: Two-Way RM ANOVA, Days, F_(1.9, 69.6)_ = 3.495; p=0.0385) after repeated cocaine exposure. However, the locomotor response to cocaine of GDX males on day 13 was not different from that on day 1 (Fig 2D: Tukey’s Post-hoc comparison (Day 1 vs Day 13, p=0.0802). These results highlight the importance of gonadal hormones in mediating neural adaptations underlying cocaine sensitization.

**Figure 2.**
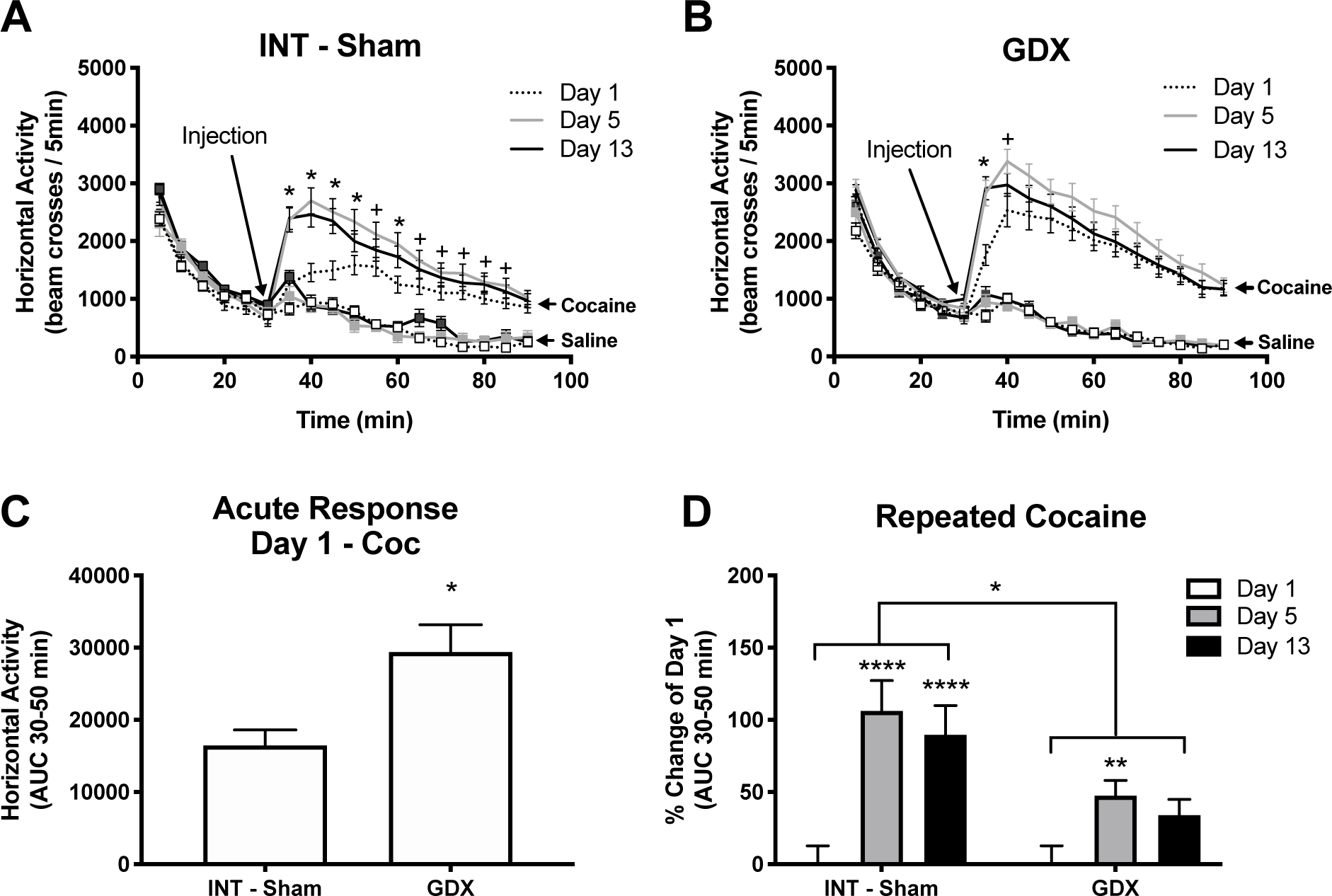
Gonadectomy increases the acute locomotor response to cocaine, and impairs sensitization in male rats. Data are represented by the average and the error bars represent the S.E.M. * p<0.05; ** p<0.01; *** p<0.001; **** P<0.0001. (See Tables 4 and 5 in the Appendix for complete statistical results). **A and B:** Time course of the locomotor activity of Intact-Sham or GDX rats injected daily with saline or with cocaine (15 mg/kg). An increase in cocaine-induced locomotor activity with time (days) was observed in intact-sham rats, and to a lesser extent, in gonadectomized (GDX), rats. The asterisks (*) represents time points in which both Day 5 and Day 13 were significantly different than Day 1 (p<0.05). The plus signs (+) represent time points in which only one of the days, Day 5 or Day 13, were significantly different than Day 1 (P<0.05). **C:** LMA represented as area under the curve (AUC) during the first 20 minutes after the first cocaine injection in intact-sham (INT-Sham) and gonadectomized (GDX) males. LMA of GDX males is significantly higher than that of INT-Sham rats **D**: Percent change of LMA of Days 5 and 13 to Day 1 of INT-Sham and GDX rats. Data represents AUC during the first 20 min after cocaine injection. INT-Sham rats showed a significant increase in cocaine-induced LMA on Days 5 and 13 compared to that displayed on Day 1, whereas GDX show an increase on Day 5, but not on Day 13.

### DHT does not restore cocaine sensitization

We demonstrated that testosterone administration enhances behavioral sensitization in GDX male rats (Menéndez-Delmestre and Segarra, 2011). To determine if this effect was mediated by testosterone’s androgenic properties, GDX animals were treated with DHT, a non-aromatizable metabolic product of testosterone, and tested for behavioral sensitization. DHT treatment had no effect on the acute locomotor response to cocaine (Fig 3B: GDX vs GDX-DHT, Tukey’s Post-hoc comparison, p=0.4767). It also did not induce cocaine sensitization in male rats (Fig 3A: Two-Way RM ANOVA, Days, F_(1.8, 39.9)_ = 0.3185; p<0.7081). Moreover, cocaine-induced LMA over time was not different between GDX and GDX-DHT (Fig 3C: Two-Way RM ANOVA, DHT X Days, F_(2, 118)_ = 0.3499; p=0.7055), but was different between INT-Sham and GDX males (Fig 3D: Two-Way RM ANOVA, DHT X Days, F_(2, 118)_ = 4.244 p=0.0166). These results suggests that the androgenic properties of testosterone do not mediate cocaine sensitization.

**Figure 3.**
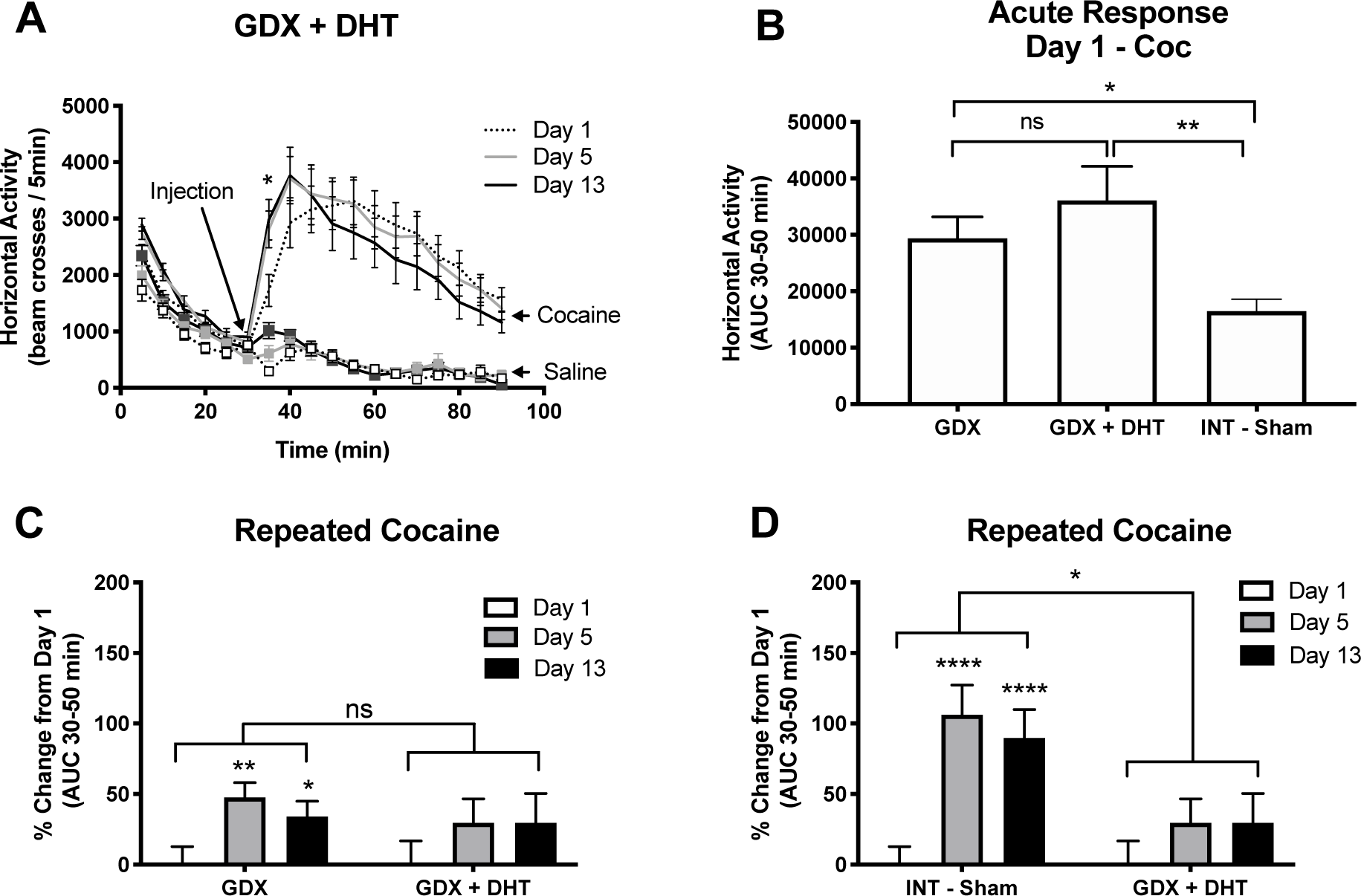
Dihydrotestosterone (DHT) has little or no effect on the acute locomotor response to cocaine and on sensitization. Data are represented by the average and the error bars represent the S.E.M. * p<0.05; ** p<0.01; *** p<0.001; **** P<0.0001. (See Tables 4 and 5 in the Appendix for complete statistical results). **A:** Time course of the locomotor activity of GDX-DHT males injected daily with saline or with cocaine (15 mg/kg). An increase in cocaine-induced locomotor activity in a single time point (5 min after cocaine administration) was observed on Days 5 and 13 compared to Day 1. The asterisks (*) represents time points in which both Day 5 and Day 13 were significantly different than day 1 (p<0.05). **B:** LMA represented as area under the curve (AUC) during the first 20 minutes after the first cocaine injection in GDX, GDX-DHT and intact-sham (INT-Sham) males. LMA of GDX and GDX-DHT males is significantly higher than that of INT-Sham rats. **C:** Percent change in LMA represented as AUC during the first 20 minutes after cocaine injection in GDX and GDX-DHT males. A RM ANOVA reveals no differences across time between GDX and GDX-DHT rats. However post hoc analysis indicate that GDX males show sensitization whereas GDX-DHT males do not (See Table 4 for complete statistical results). This issue is further analyzed in the discussion section. **D:** Percent change in LMA represented as AUC during the first 20 minutes after cocaine injection in INT-Sham and GDX-DHT males. A RM ANOVA reveals a significant difference across time between INT-Sham and GDX-DHT rats, INT-Sham rats display sensitization whereas GDX-DHT do not.

### Estradiol restores cocaine sensitization

Testosterone can also be aromatized to estradiol. To investigate whether testosterone’s enhancement of cocaine sensitization in male rats is mediated through estradiol we treated GDX animals with EB systemically and tested for behavioral sensitization. EB treatment did not affect the acute locomotor response to cocaine (Fig 4B: GDX vs GDX-EB, Tukey’s Post-hoc comparison, p=0.9700). However, EB treated GDX males, similar to INT-Shams, displayed a robust sensitization (Fig 4A:Two-Way RM ANOVA, Days, F_(1.9, 42.6)_ = 18.79; p<0.0001; Fig 4D: Two-Way RM ANOVA, GDX-EB X Days, F_(2, 118)_ = 1.134; p=0.3252). This robust sensitization was significantly different from that displayed by GDX males (Fig 4C: Two-Way RM ANOVA, EB X Days, F_(59,118)_ = 2.838; p<0.0001). These results show that estradiol modulates sensitization to cocaine in GDX males, and suggest that estradiol is important for male rats to become sensitized to cocaine.

**Figure 4.**
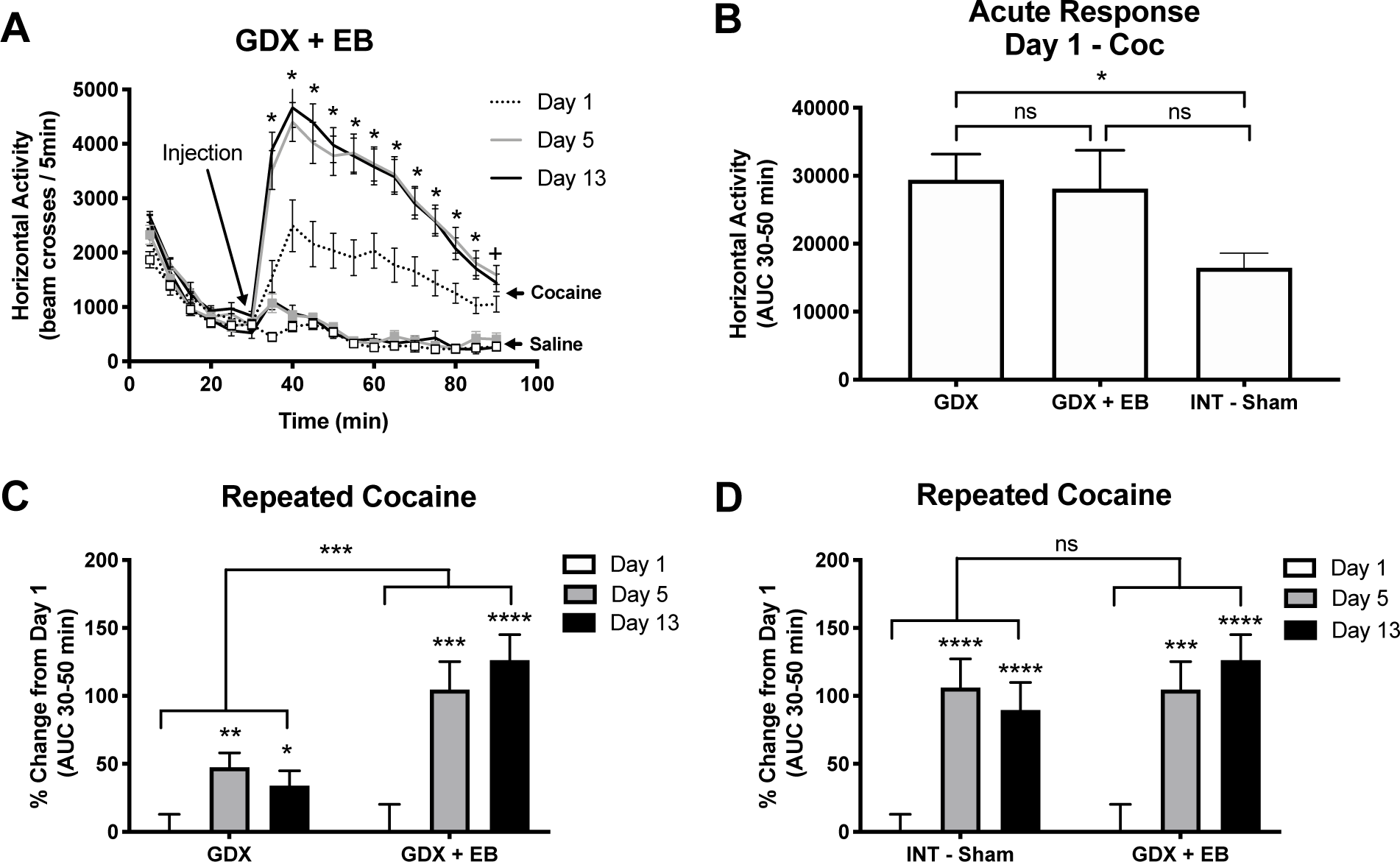
Locomotor response to cocaine of gonadectomized male rats treated with estradiol. Data are represented by the average and the error bars represent the S.E.M. * p<0.05; ** p<0.01; *** p<0.001; **** P<0.0001. (See Tables 4 and 5 in the Appendix for complete statistical results). **A:** Time course of the locomotor activity of GDX estradiol-treated (EB) males injected daily with saline or with cocaine (15 mg/kg). An increase in cocaine-induced locomotor activity at all time points was observed on days 5 and 13 (except min 90) compared to Day 1. The asterisks (*) represents time points in which both Day 5 and Day 13 were significantly different than Day 1 (p<0.05). The plus sign (+) represents time points in which only one of the days, Day 5 or Day 13, was significantly different than Day 1 (P<0.05). **B:** LMA represented as area under the curve (AUC) during the first 20 minutes after the first cocaine injection in GDX, GDX-EB and intact-sham (INT-Sham) males. LMA of GDX and GDX-EB males is higher than that of INT-Sham rats, but this difference reached significance only in GDX males. **C:** Percent change in LMA represented as AUC during the first 20 minutes after cocaine injection in GDX and GDX-EB males. A RM ANOVA reveals dramatic differences across time between GDX and GDX-EB rats, indicating a more robust sensitization in GDX-EB rats. **D:** Percent change in LMA represented as AUC during the first 20 minutes after cocaine injection in INT-Sham and GDX-EB males. A RM ANOVA reveals no differences across time between INT-Sham and GDX-EB rats, indicating that estrogen restores sensitization to that of INT-Sham animals.

### Estrogen receptor inhibition blocks cocaine sensitization

To determine the role of estrogen receptors in cocaine-induced locomotor activity we blocked central ERs with ICI 182,780. Repeated cocaine administration resulted in behavioral sensitization of gonadally intact control males (INT+Veh) (Fig 5A: Two-Way RM ANOVA, Days, F_(1.2, 9.6)_ = 8.343 p=0.0139), an effect blocked by ICI 182,780 treatment (INT+ICI) (Fig 5B: Two-Way RM ANOVA, Days, F_(1.2, 7.0)_ = 1.986; p=0.2043). These results indicate that gonadally-intact males require activation of central estrogen receptors for the expression of robust behavioral sensitization to cocaine.

**Figure 5.**
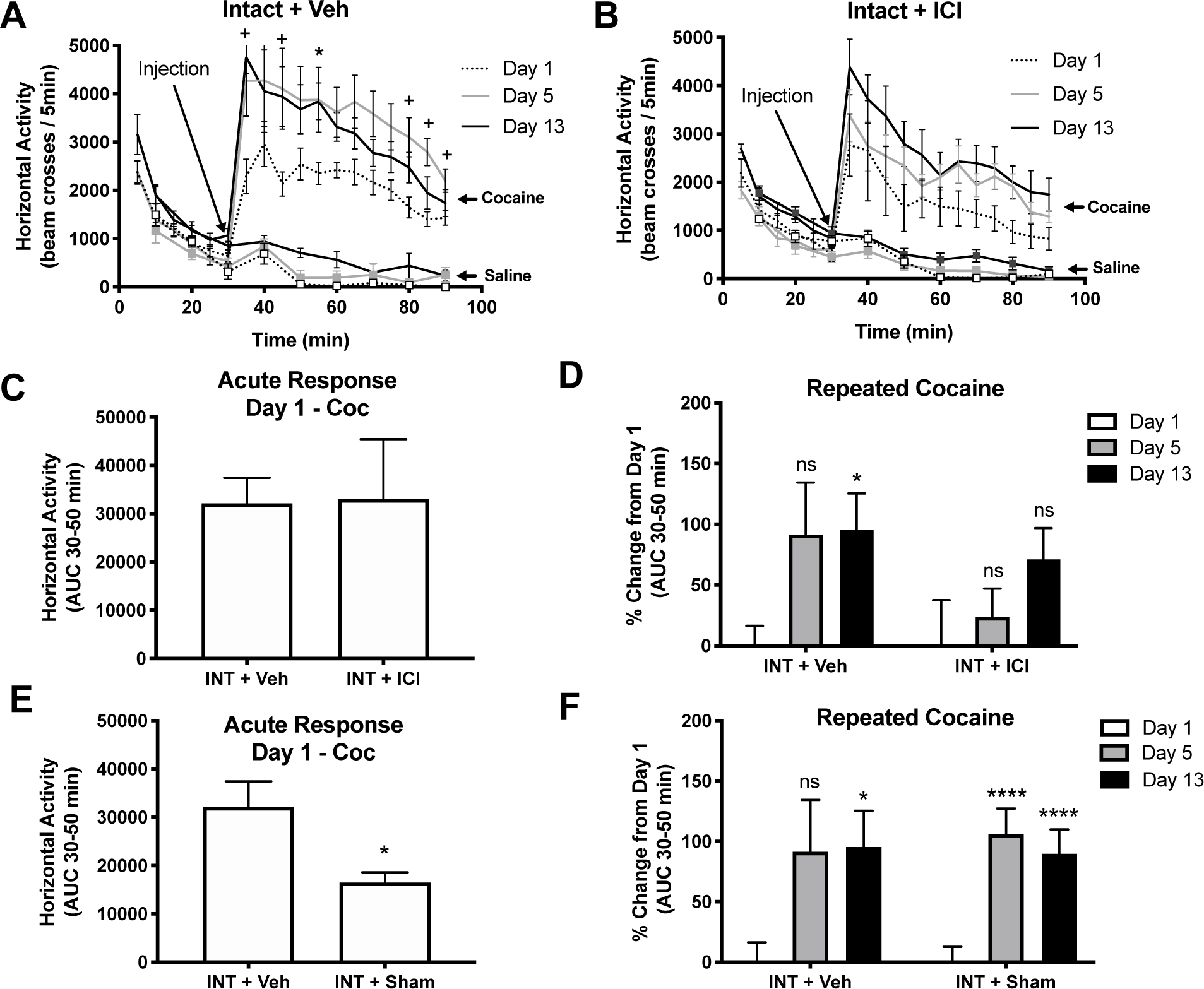
The antiestrogen ICI-182,780 blocks sensitization to cocaine in intact male rats. Data are represented by the average and the error bars represent the S.E.M. * p<0.05; ** p<0.01; *** p<0.001; **** P<0.0001. (See Tables 4 and 5 in the Appendix for complete statistical results). **A and B:** Time course of the locomotor activity of intact-vehicle (INT-Veh) or INT-ICI rats injected daily with saline or with cocaine (15 mg/kg). An increase in cocaine-induced locomotor activity with time (days) was observed in INT-Veh, and not in rats treated with ICI 182,780. The asterisks (*) represents time points in which both Day 5 and Day 13 were significantly different than Day 1 (p<0.05). The plus signs (+) represent time points in which only one of the days, Day 5 or Day 13, were significantly different than Day 1 (P<0.05). **C:** LMA represented as area under the curve (AUC) during the first 20 minutes after the first cocaine injection in INT-Veh and INT-ICI males. No difference was observed in the acute locomotor response to cocaine between these groups. **D:** Percent change in LMA represented as AUC during the first 20 minutes after cocaine injection in INT-Veh and INT-ICI males. A RM ANOVA reveals no differences across time between INT-Veh and INT-ICI rats. However post hoc analysis indicate that INT-Veh show sensitization that reached significance by day 13 whereas INT-ICI males do not sensitize to cocaine. **E:** LMA represented as area under the curve (AUC) during the first 20 minutes after the first cocaine injection in INT-Veh and INT-Sham males. INT-Veh males show a higher locomotor response to cocaine on Day 1 than INT-Sham rats. **F:** Percent change in LMA represented as AUC during the first 20 minutes after cocaine injection in INT-Veh and INT-Sham males. A RM ANOVA reveals no differences across time between INT-Veh and INT-ICI rats. However post hoc analysis indicate that INT-Veh show sensitization that reached significance by Day 13 whereas sensitization in INT-Sham males is evident on Days 5 and 13.

To examine if ICI has an effect on basal locomotor activity, we compared basal LMA of INT+ICI saline males to that of INT+Veh saline males each day tested (Days 1, 5 and 13) using a Repeated Measures ANOVA. The data analysis shows that LMA did not differ among saline males in any of the days tested (F_(1, 14)_ = 0.7467; p=0.4021).

Interestingly, cocaine-induced LMA on day 1 did not differ between INT+Veh and INT-ICI males (Fig 5C: Student’s T-test, t=0.06286, df= 8.165, p=0.9514). Further perusal of the results show that INT-Veh males display higher cocaine-induced LMA on day 1 than INT-Sham males (Fig 5E: Student’s T-test, t=2.754, df= 10.69, p=0.0192), but both groups display sensitization to cocaine (Fig 5F: Two-Way RM ANOVA, Sham-Veh X Days, F_(2, 90)_ = 0.1363, p=0.8728). INT-Sham males were sham operated for testes whereas INT-Veh rats were subjected to stereotaxic surgery and implanted with an Alzet mini-pump that delivered vehicle into the lateral ventricle. This finding suggests that the procedure of stereotaxic surgery exacerbates the acute locomotor response to cocaine. These data highlight the complexity of locomotor activity and the importance of appropriate controls.

## Discussion

We had previously demonstrated that gonadectomy increased the acute locomotor response to cocaine but prevented the progressive increase in locomotion over time observed after repeated injections, i.e. cocaine-induced behavioral sensitization (Menéndez-Delmestre and Segarra, 2011). We had also reported that testosterone administration to GDX rats restored sensitization in male rats and decreased the acute locomotor response to cocaine (Menéndez-Delmestre and Segarra, 2011). The present study extends these findings and reports that estradiol, but not DHT, enhances the locomotor response to repeated cocaine administration. These results suggest that in male rats, androgen receptors do not participate in mediating neuroadaptations resulting from long-term drug administration. Furthermore, we report that intracellular ERs mediate the enhancement of cocaine-induced locomotor activity over time. This study found that blocking central ERs eliminates cocaine-induced behavioral sensitization. To our knowledge, this is the first study that illustrates the pivotal role of ERs in cocaine sensitization in males, suggesting that estrogen receptors play a key role in cocaine-induced neuroadaptations. Moreover, this study indicates that estrogen signaling participates in mediating drug-related behaviors in males and is also a key signal of cocaine-induced neuroplasticity in the rodent brain.

### Gonadal steroids and the acute response to cocaine

Our data indicate that gonadectomy increases the locomotor response to cocaine in drug naive rats (i.e. day 1). Neither estradiol nor DHT replacement were sufficient to prevent this increase, however testosterone replacement was (Menendez-Delmestre and Segarra, 2011). Given that testosterone is normally transformed into DHT and estradiol, our combined findings suggest that activation of both receptors (AR and ER) play a role in regulating the psychostimulant response to cocaine in drug naive animals.

### Gonadal steroids and sensitization to cocaine

Even though sensitization is more robust in gonadally intact males, and in those that receive estradiol replacement, GDX males do sensitize as evidenced by the percent change in locomotor activity between days 1 and days 5 (Fig 2D). These results differ from those we published in our previous paper (Menéndez-Delmestre and Segarra, 2011). We believe that an increase in the number of animals in this study can partially explain these differences. We encountered the same situation with our studies of ovariectomized female rats (Segarra et al., 2014) where increasing the number of animals turned a consistent trend into significantly different results.

This study found that DHT did not affect the acute nor enhanced the sensitized response to cocaine. On the contrary, DHT would appear to block sensitization (Fig 3C). However, we are cautious to embrace this finding since our statistical analysis show conflicting results. For instance, when comparing the timecourse of GDX and GDX-DHT (Figs 2B and 3A), although both groups show a significant difference in cocaine-induced LMA from day 1 in the first 5 min after cocaine injection (min 35), GDX males also show a difference at timepoint 40 (Day1 vs Day 5). In addition, post-hoc analysis of Fig 3C (2-way ANOVA-Table 4) indicates that GDX males sensitize whereas GDX-DHT do not. While these two findings are consistent with the idea that DHT suppresses sensitization, a trend of higher locomotor activity is also observed in GDX-DHT males on days 5 and 13. Moreover, the interaction of the overall Two Way ANOVA indicates there are no significant differences in the locomotor response to cocaine over time between GDX and GDX-DHT males (see Table 4 of Appendix). Accordingly, further studies are required to elucidate the role of DHT on cocaine sensitization in male rodents.

Estradiol is the gonadal hormone that appears to play a greater role in mediating cocaine-induced neuroadaptations after repeated cocaine administration, since T and EB are capable of sustaining behavioral sensitization in gonadectomized animals, whereas DHT is not. Blocking ER in intact male rats with ICI does not alter the locomotor response to cocaine on day 1 but blocks sensitization to cocaine as well. Moreover, our findings with the antiestrogen ICI suggests that central ER, play a key role in mediating cocaine-induced neuroadaptations after repeated cocaine administration. We do not believe that a ceiling effect is responsible for the lower sensitization displayed by GDX males since there are several instances where animals exhibit a further increase in their locomotor response to cocaine, even though their response on day 1 was high. For instance, the response of GDX males on day 1 is similar to that of GDX-EB rats (Fig 2B), however, sensitization of GDX-EB is very robust (Fig 4C). Additionally, in our previous study, GDX males show further increases in locomotor activity with higher doses of cocaine (Menéndez-Delmestre and Segarra, 2011), which is inconsistent with a ceiling effect.

Gonadectomy has broader consequences than the selective elimination of gonadal hormones, such as alteration of hypothalamic and hypophyseal hormone levels (Gay et al., 1970; Hood and Schwartz, 2000; Jansson et al., 1984), and changes in neuroplasticity (Cooke and Woolley, 2009; Harley et al., 2000; Hebbard et al., 2003; Leranth et al., 2003; Spencer et al., 2008). Even though many of these effects can be attributed to changes in plasma gonadal steroid levels, exogenous hormonal administration does not restore the pattern, or cyclicity of secretion. Differences in hormonal replacement methods, which as a consequence produce variations in the plasma hormonal levels attained, can also alter the results obtained. To overcome this caveat we used the anti-estrogen ICI-182,780 administered directly into the right lateral ventricle of the brain to avoid the controversy of whether the compound can cross the blood brain barrier (Alfinito et al., 2008; Dehghan et al., 2015; Wakeling et al., 1991). Our findings with the estrogen receptor antagonist (ICI-182,780) in intact rats (INT-ICI) confirm the importance of ER activation in cocaine-induced neuroplasticity of male rats.

The role of estradiol in cocaine-induced behavioral sensitization in females has been extensively characterized (Segarra et al., 2014). Cocaine-induced hyperactivity varies throughout the estrous cycle, being highest at estrous, just after the estradiol surge (Quiñones-Jenab et al., 1999; Sell et al., 2002, 2000). In addition, estradiol administration to OVX female rats restores (Febo et al., 2003; Hu and Becker, 2003; Puig-Ramos et al., 2008; Segarra et al., 2010) and increases sensitization to cocaine as well as conditioned place preference (Segarra et al., 2014). Treatment of gonadally intact rats (INT+ICI) with the estrogen receptor antagonist ICI-182,780 confirm the importance of ER activation in cocaine-induced neuroplasticity (Segarra et al., 2014). New studies also highlight the importance of estradiol in mediating amphetamine stimulated DA release in the dorsolateral striatum of female rats (Song et al., 2019). The nucleus accumbens participates in the regulation of locomotor activity and incentive motivational behaviors, and contains estrogen receptors. Indeed, estradiol increases medium spiny neuron dendritic density, changes striatal electrophysiological properties (Meitzen et al., 2018) and increase extracellular dopamine (for review see Segarra et al., 2010).

Recent studies are uncovering the multiple effects of estradiol in males. Treatment with estradiol of aromatase deficient man highlight the importance of estradiol in the regulation of metabolic and hepatic function (for review see Song et al., 2019; Meitzen et al., 2018; Santen and Simpson, 2019). Estradiol enhances insulin sensitivity and glucose uptake (Díaz et al., 2019), semen quality (Yuan et al., 2019), enhances long term memory (Mitchnik et al., 2019), and exerts neuroprotective effects (Chen et al., 2019). Indeed, there are reported instances where the effects of testosterone are exerted by interacting with estrogen receptors. For example, the neuroprotective properties of testosterone against the neurotoxic effects of the HIV proteins gp120 and Tat is blocked by the anti-estrogen ICI-182,780 (Kendall et al., 2005). Furthermore, several studies demonstrate that in the male, estradiol and estrogen receptors also play a significant role in the regulation of neural processes and behaviors. In a male model of Parkinson disease, administration of ICI-182,780 blocked the protective effect of estradiol on dopaminergic neurons (Bourque et al., 2015). In male mice, an increase in anxiety and in GAD 65 was found after intracerebroventricular administration of letrozole, an aromatase inhibitor (Ikeda et al., 2015). Studies by Mitchnik et al. (2019) demonstrate the importance of ERs in perirhinal cortex-mediated short term and long term object-in-place memory in male rats. Thus, the notion of male and female gonadal hormones is currently eroding as recent articles emphasize the contribution of estradiol in males and of testosterone in females, in the maintenance of basic physiological processes (Hammes and Levin, 2019; Russell and Grossmann, 2019).

ERs, alpha and beta, are expressed in dopaminergic neurons of the VTA (Creutz and Kritzer, 2004; Kritzer, 1997; Kritzer and Creutz, 2008), a central component of the reward circuit that projects to limbic and forebrain areas, including the nucleus accumbens (NAc) and prefrontal cortex (PFC). Activity of VTA dopaminergic neurons is regulated, in part, by VTA GABAergic tone. Estradiol inactivates GABA_B_ receptors (Kelly et al., 2003; Qiu et al., 2008; Roepke et al., 2009), thus increasing neuronal firing and neurotransmitter release from dopaminergic neurons in the VTA. We have also shown that systemic estradiol administration to OVX rats decreases GABA_B_ activity in the VTA (Febo and Segarra, 2004). Extrapolating these data to our behavioral studies we hypothesize that intact (INT+Veh, and INT-Sham) and GDX-EB animals increase their behavioral response to repeated cocaine due to increased activation of ERs in the VTA. This would lead to decreased inhibition on dopaminergic neurons, resulting in increased DA release into the NAc and cocaine sensitization. In contrast, blocking ERs with ICI-182,780 would increase inhibition on dopaminergic neurons, resulting in suppression of behavioral sensitization to cocaine. However, GABA_B_ activation in the VTA of male rats has not been examined in this study and requires further investigation.

Repeated cocaine administration also results in strengthening of glutamatergic synapses, increasing sensitivity to excitatory inputs (Koob and Volkow, 2010; Riegel and Kalivas, 2010). Estradiol has been shown to act both acutely, by membrane-initiated signaling of the ER, and chronically, by nuclear-initiated signaling of the ER. Acute activation of ERs activates cAMP/PKA signaling pathway that is hypothesized to produce cAMP-dependent phosphorylation events (Kelly and Rønnekleiv, 2008). Nuclear-initiated signaling activates the intracellular ER, which has been shown to upregulate transcription of kinases, phosphatases, signaling and transcriptional factors that could explain the robust sensitized behavioral response of estradiol treated GDX males to repeated cocaine administration (for review see Kelly et al., 2003).

In summary, this study extends our previous findings that highlight the importance of testosterone in male rats to become sensitized to cocaine. The current study suggests that in males, testosterone aromatizes to estradiol and activates ERs and in this way enhances sensitization to cocaine. We propose that ERs are the biological substrates through which gonadal steroids enhance sensitization to cocaine, irrespective of sex. Further studies are required to elucidate the molecular mechanisms involved in estradiol modulation of cocaine-sensitization in male and female rats. In addition, these data show the importance of evaluating the hormonal status of individuals prior and during treatment for cocaine addiction.

## Acknowledgements

The authors would like to thank Natasha Lugo-Escobar, Anabel Puig-Ramos, Rafael Vázquez, María Carolina Velázquez, María Vélez, Waldo Amadeo, Richard D. Silva and José G. Rivera for their technical assistance. This work was supported by the National Science Foundation [grants numbers: PIRE 1545803, IIS 1633184] and from the National Institutes of Health [RR03051(RCMI); U54NS39405 (SNRP); and R25GM061838 (MBRS/RISE)].

---

**Table A1:** Number of animals in each treatment group.

| Table A1: Number of animals in each treatment group |  |  |  |  |
| --- | --- | --- | --- | --- |
| Treatment | Total # of animals (Saline) | Total # of animals (Cocaine) | Number of experiments | Number of animals/exp |
| Intact-Sham | 18 | 38 | 12 | 2-5 |
| GDX | 20 | 38 | 12 | 2-5 |
| GDX-DHT | 18 | 23 | 4 | 4-7 |
| GDX-EB | 18 | 23 | 4 | 5-7 |
| INT-Veh | 5 | 9 | 1 | N/A |
| INT-ICI | 8 | 7 | 1 | N/A |

**Table A2:**
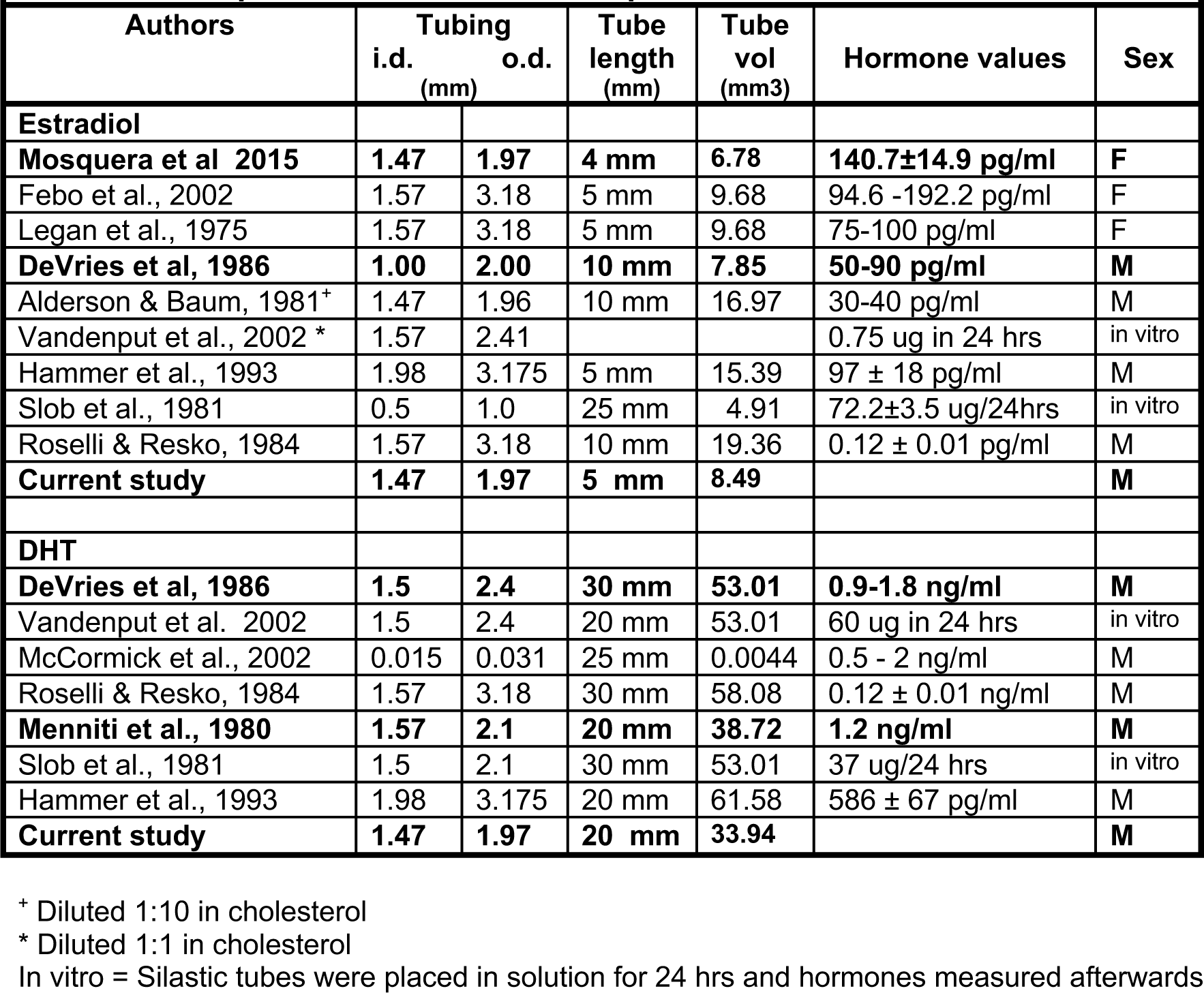
Comparison of Silastic tube implants and hormonal levels achieved.

**Table A3.** Summary of the Normality tests of all data included in Figures 2-5.

| Treatment | Timepoint (Day) | Measurement Unit | Sample Size (N) | Test for normal distribution D'Agostino & Pearson test |  |  | Corresponding Figures |
| --- | --- | --- | --- | --- | --- | --- | --- |
|  |  |  |  | K2 | P-Value | Passed Norm. Test |  |
| INT + Sham | 1 | % change of day 1 | 38 | 5.352 | 0.0688 | Yes | Figs. 2D,3D,4D,5F |
|  | 5 |  | 38 | 4.99 | 0.0825 | Yes |  |
|  | 13 |  | 38 | 4.312 | 0.1158 | Yes |  |
| GDX | 1 | % change of day 1 | 38 | 10.07 | 0.0065 | No | Figs. 2D,3C,4C |
|  | 5 |  | 38 | 1.651 | 0.438 | Yes |  |
|  | 13 |  | 38 | 2.393 | 0.3023 | Yes |  |
| GDX + DHT | 1 | % change of day 1 | 23 | 6.995 | 0.0303 | No | Figs. 3C, 3D |
|  | 5 |  | 23 | 3.256 | 0.1964 | Yes |  |
|  | 13 |  | 23 | 4.54 | 0.1033 | Yes |  |
| GDX + EB | 1 | % change of day 1 | 23 | 3.857 | 0.1454 | Yes | Figs. 4C, 4D |
|  | 5 |  | 23 | 0.003327 | 0.9983 | Yes |  |
|  | 13 |  | 23 | 4.244 | 0.1198 | Yes |  |
| INT + Veh | 1 | % change of day 1 | 9 | 1.103 | 0.5761 | Yes | Figs. 5D, 5F |
|  | 5 |  | 9 | 2.101 | 0.3498 | Yes |  |
|  | 13 |  | 9 | 1.934 | 0.3802 | Yes |  |
| INT + ICI | 1 | % change of day 1 | 7 | N too small | 0.0222* | No | Fig. 5D |
|  | 5 |  | 7 | N too small | 0.3212* | Yes |  |
|  | 13 |  | 7 | N too small | 0.8408* | Yes |  |

**Table A4.**
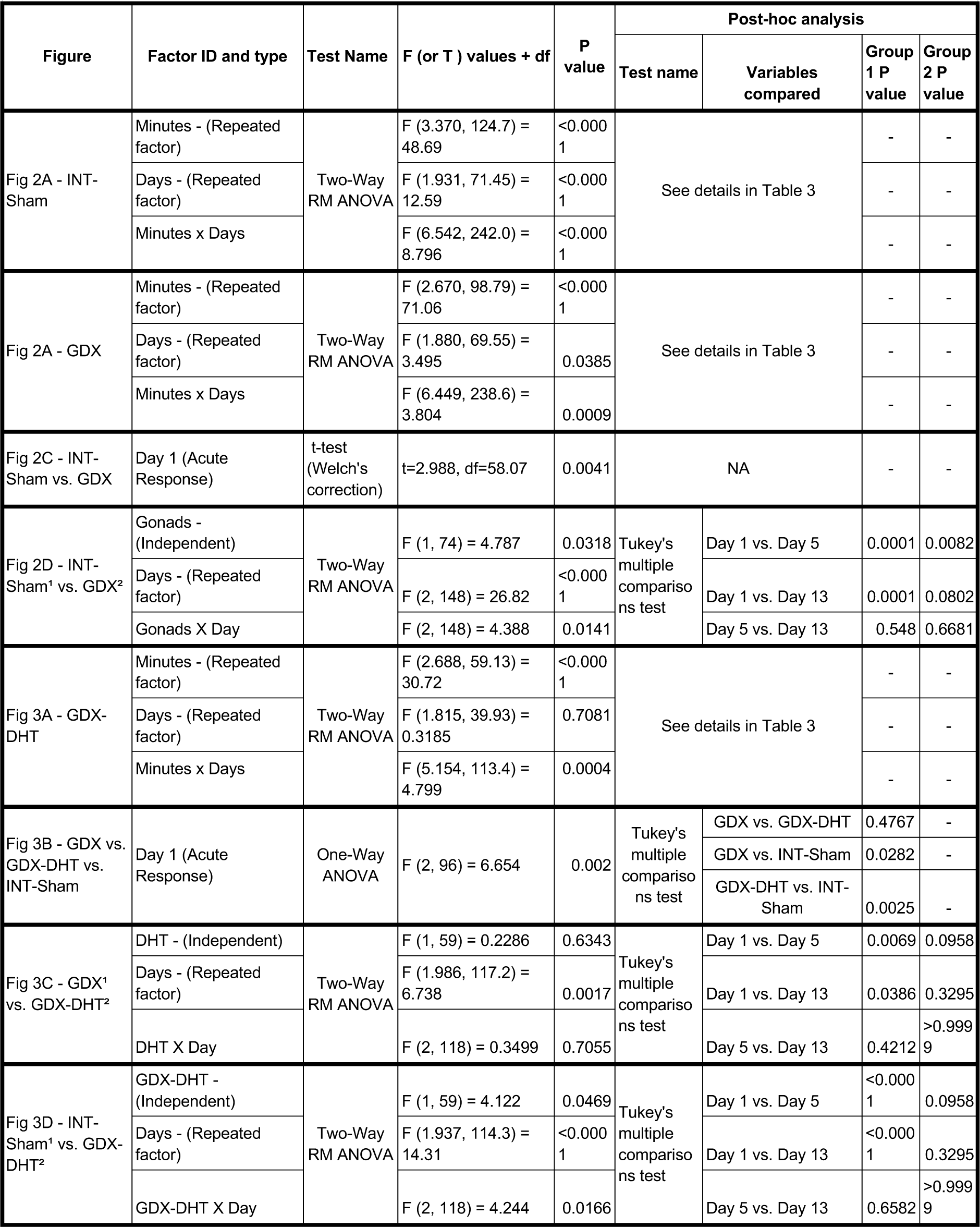
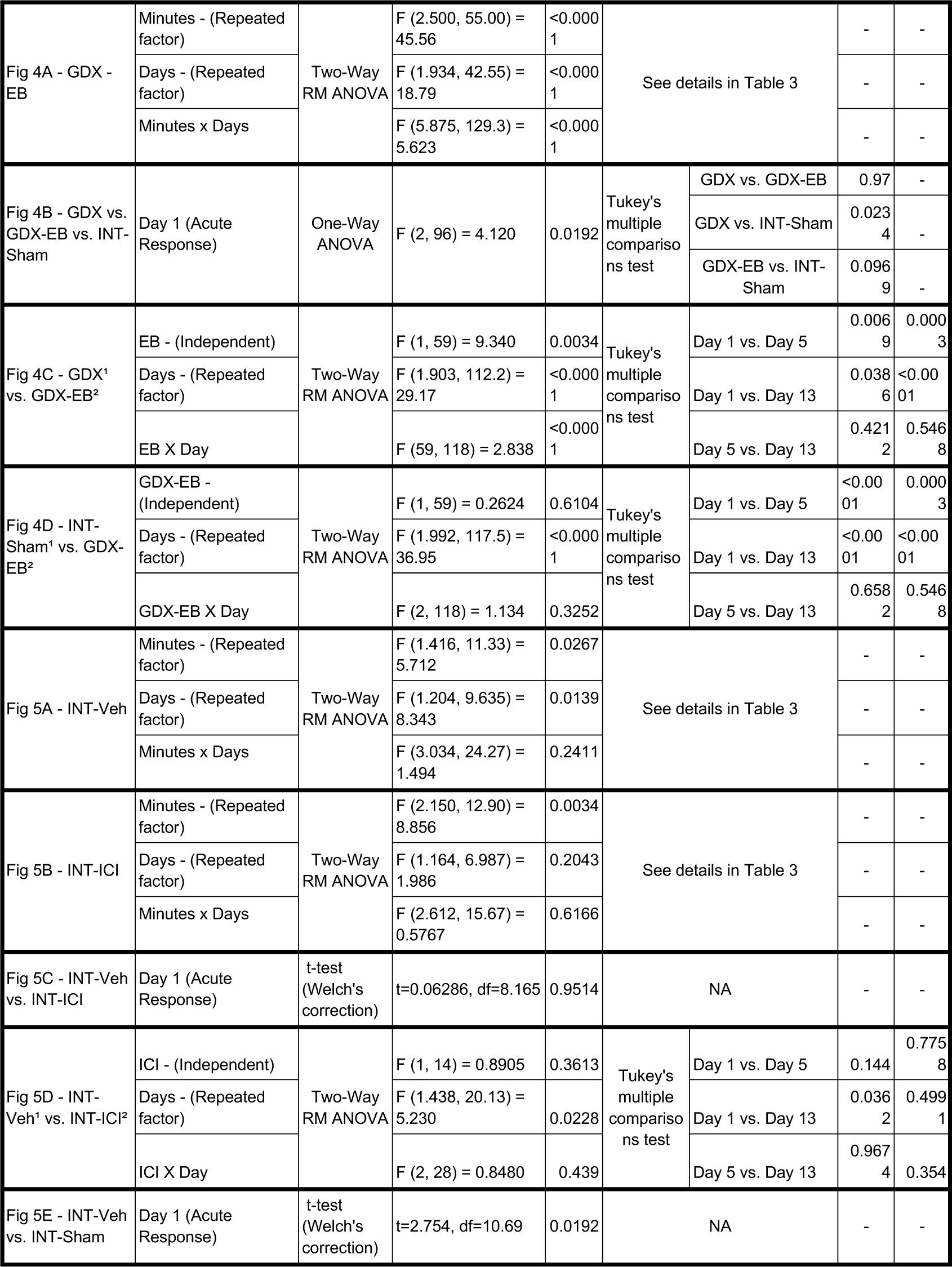
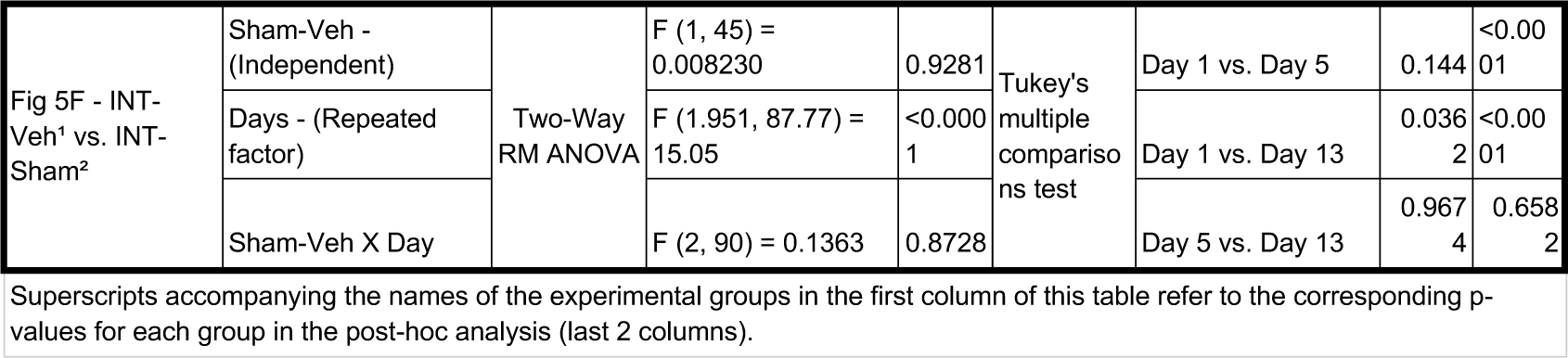
Statistical table for Figures 2-5.

**Table A5.**
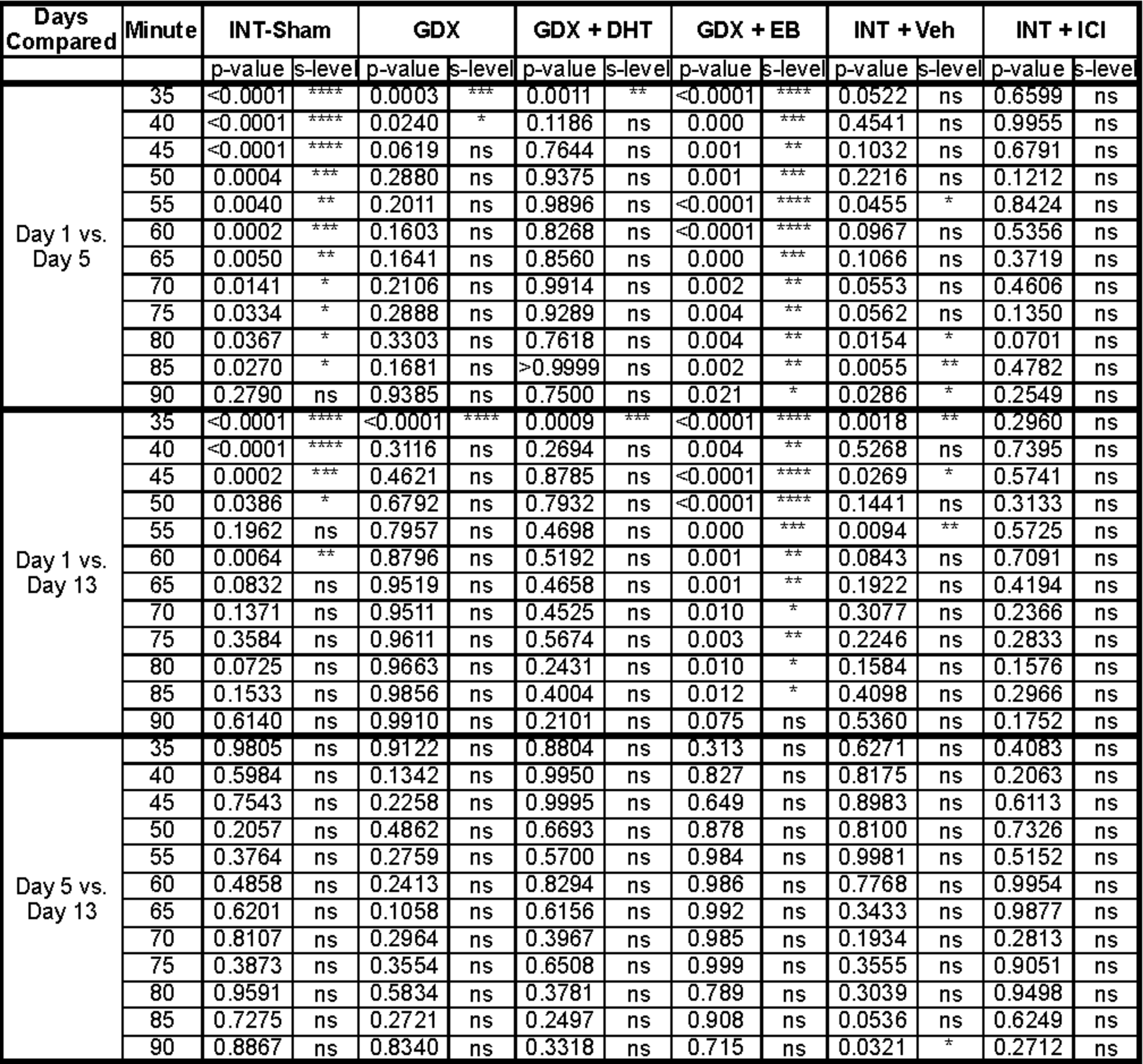
Post-hoc analysis of the time course of each experimental group using Tukey’s multiple comparisons.

| Days Compared | Minute | INT-Sham |  | GDX |  | GDX + DHT |  | GDX + EB |  | INT + Veh |  | INT + ICI |  |
| --- | --- | --- | --- | --- | --- | --- | --- | --- | --- | --- | --- | --- | --- |
|  |  | p-value | s-level | p-value | s-level | p-value | s-level | p-value | s-level | p-value | s-level | p-value | s-level |
| Day 1 vs. Day 5 | 35 | <0.0001 | **** | 0.0003 | *** | 0.0011 | ** | <0.0001 | **** | 0.0522 | ns | 0.6599 | ns |
|  | 40 | <0.0001 | **** | 0.0240 | * | 0.1186 | ns | 0.000 | *** | 0.4541 | ns | 0.9955 | ns |
|  | 45 | <0.0001 | **** | 0.0619 | ns | 0.7644 | ns | 0.001 | ** | 0.1032 | ns | 0.6791 | ns |
|  | 50 | 0.0004 | *** | 0.2880 | ns | 0.9375 | ns | 0.001 | *** | 0.2216 | ns | 0.1212 | ns |
|  | 55 | 0.0040 | ** | 0.2011 | ns | 0.9896 | ns | <0.0001 | **** | 0.0455 | * | 0.8424 | ns |
|  | 60 | 0.0002 | *** | 0.1603 | ns | 0.8268 | ns | <0.0001 | **** | 0.0967 | ns | 0.5356 | ns |
|  | 65 | 0.0050 | ** | 0.1641 | ns | 0.8560 | ns | 0.000 | *** | 0.1066 | ns | 0.3719 | ns |
|  | 70 | 0.0141 | * | 0.2106 | ns | 0.9914 | ns | 0.002 | ** | 0.0553 | ns | 0.4606 | ns |
|  | 75 | 0.0334 | * | 0.2888 | ns | 0.9289 | ns | 0.004 | ** | 0.0562 | ns | 0.1350 | ns |
|  | 80 | 0.0367 | * | 0.3303 | ns | 0.7618 | ns | 0.004 | ** | 0.0154 | * | 0.0701 | ns |
|  | 85 | 0.0270 | * | 0.1681 | ns | >0.9999 | ns | 0.002 | ** | 0.0055 | ** | 0.4782 | ns |
|  | 90 | 0.2790 | ns | 0.9385 | ns | 0.7500 | ns | 0.021 | * | 0.0286 | * | 0.2549 | ns |
| Day 1 vs. Day 13 | 35 | <0.0001 | **** | <0.0001 | **** | 0.0009 | *** | <0.0001 | **** | 0.0018 | ** | 0.2960 | ns |
|  | 40 | <0.0001 | **** | 0.3116 | ns | 0.2694 | ns | 0.004 | ** | 0.5268 | ns | 0.7395 | ns |
|  | 45 | 0.0002 | *** | 0.4621 | ns | 0.8785 | ns | <0.0001 | **** | 0.0269 | * | 0.5741 | ns |
|  | 50 | 0.0386 | * | 0.6792 | ns | 0.7932 | ns | <0.0001 | **** | 0.1441 | ns | 0.3133 | ns |
|  | 55 | 0.1962 | ns | 0.7957 | ns | 0.4698 | ns | 0.000 | *** | 0.0094 | ** | 0.5725 | ns |
|  | 60 | 0.0064 | ** | 0.8796 | ns | 0.5192 | ns | 0.001 | ** | 0.0843 | ns | 0.7091 | ns |
|  | 65 | 0.0832 | ns | 0.9519 | ns | 0.4658 | ns | 0.001 | ** | 0.1922 | ns | 0.4194 | ns |
|  | 70 | 0.1371 | ns | 0.9511 | ns | 0.4525 | ns | 0.010 | * | 0.3077 | ns | 0.2366 | ns |
|  | 75 | 0.3584 | ns | 0.9611 | ns | 0.5674 | ns | 0.003 | ** | 0.2246 | ns | 0.2833 | ns |
|  | 80 | 0.0725 | ns | 0.9663 | ns | 0.2431 | ns | 0.010 | * | 0.1584 | ns | 0.1576 | ns |
|  | 85 | 0.1533 | ns | 0.9856 | ns | 0.4004 | ns | 0.012 | * | 0.4098 | ns | 0.2966 | ns |
|  | 90 | 0.6140 | ns | 0.9910 | ns | 0.2101 | ns | 0.075 | ns | 0.5360 | ns | 0.1752 | ns |
| Day 5 vs. Day 13 | 35 | 0.9805 | ns | 0.9122 | ns | 0.8804 | ns | 0.313 | ns | 0.6271 | ns | 0.4083 | ns |
|  | 40 | 0.5984 | ns | 0.1342 | ns | 0.9950 | ns | 0.827 | ns | 0.8175 | ns | 0.2063 | ns |
|  | 45 | 0.7543 | ns | 0.2258 | ns | 0.9995 | ns | 0.649 | ns | 0.8983 | ns | 0.6113 | ns |
|  | 50 | 0.2057 | ns | 0.4862 | ns | 0.6693 | ns | 0.878 | ns | 0.8100 | ns | 0.7326 | ns |
|  | 55 | 0.3764 | ns | 0.2759 | ns | 0.5700 | ns | 0.984 | ns | 0.9981 | ns | 0.5152 | ns |
|  | 60 | 0.4858 | ns | 0.2413 | ns | 0.8294 | ns | 0.986 | ns | 0.7768 | ns | 0.9954 | ns |
|  | 65 | 0.6201 | ns | 0.1058 | ns | 0.6156 | ns | 0.992 | ns | 0.3433 | ns | 0.9877 | ns |
|  | 70 | 0.8107 | ns | 0.2964 | ns | 0.3967 | ns | 0.985 | ns | 0.1934 | ns | 0.2813 | ns |
|  | 75 | 0.3873 | ns | 0.3554 | ns | 0.6508 | ns | 0.999 | ns | 0.3555 | ns | 0.9051 | ns |
|  | 80 | 0.9591 | ns | 0.5834 | ns | 0.3781 | ns | 0.789 | ns | 0.3039 | ns | 0.9498 | ns |
|  | 85 | 0.7275 | ns | 0.2721 | ns | 0.2497 | ns | 0.908 | ns | 0.0536 | ns | 0.6249 | ns |
|  | 90 | 0.8867 | ns | 0.8340 | ns | 0.3318 | ns | 0.715 | ns | 0.0321 | * | 0.2712 | ns |

